# Induced Droplet Ovalisation (IDO): Image-based microfluidic method for high-throughput and label-free characterization of microbial proteolytic strains from wastewater sludge

**DOI:** 10.1101/2025.05.28.656601

**Authors:** Luca Potenza, Maciej S. Andrzejewski, Tomasz S. Kaminski

## Abstract

Traditional bacterial isolation methods are costly, labour-intensive, and often fail to detect rare or slow-growing taxa. In contrast, droplet-based microfluidic assays have emerged as powerful alternatives. Here, we describe a novel, label-free microfluidic assay for screening microbial proteolytic activity based on oil-induced droplet ovalization. This method outperformed two traditional isolation techniques in terms of performance, time, and overall cost-efficiency. We present Induced Droplet Ovalisation (IDO): a droplet-based protocol that combines single-cell encapsulation with automated image analysis. Using a custom-made microfluidic device and bespoke image-processing scripts, our system efficiently detects microbial proteolytic activity by monitoring droplet deformability. This approach significantly reduces the time and resources required to study proteolytic consortia, offering an automated, easy-to-implement, and label-free alternative to conventional screening. Our device achieves high-throughput screening (HTS) at 0.2 kHz while significantly reducing the cultivation medium volume used compared to traditional methods. Furthermore, in-droplet microbial recovery is enhanced by two orders of magnitude, enabling the enumeration of significantly more colonies compared to traditional screenings. Beyond microbiological applications, this versatile platform serves as a powerful tool for studying hydrogel degradation and polymerization dynamics, extending its applicability across research and industry.

## Introduction

Droplet microfluidics has also emerged as a transformative tool in high-throughput screening (HTS) of biochemical reactions, offering a promising alternative to traditional methods in dishes, flasks and multiwell plates. Droplet-based microfluidics systems have also enabled the development of novel protocols for screening microbial activity, offering key advantages such as miniaturized volumes, high throughput, and the ability to study both single cells and microbial consortia^1^. Despite these benefits, several challenges remain, including droplet stability, growth condition optimization, scalability, and strain recovery for upscaling and downstream applications. Nevertheless, these assays outperform traditional protocols in throughput, accuracy and cost efficiency, making them a powerful tool for biocatalytic screening and discovery.

Droplet microfluidic systems have been successfully applied to enrich microbes based on growth, antibiotic production, and biocatalytic activity^2–4^. Such screening assays based on fluorescence^5–7^ and absorbance^8,9^ detection have been extensively used in microfluidics to detect microbial activity, leveraging decades of established protocols and technologies. Optical systems demonstrate high sensitivity, particularly fluorescence activated droplet sorting (FADS)^5^ and Absorbance-Activated Droplet Sorting (AADS)^8^, and offer throughput with sorting capabilities exceeding kHz frequencies. However, optical assays require dyes and/or labels, which pose a significant bottleneck due to the limited number of reactions that can be implemented in a droplet format. In addition, detection setups for absorbance or fluorescence entail substantial upfront investments, including specialised light sources, detectors, and optoelectronic components.

From a biological perspective, native substrates are typically unlabeled, and introducing chemically modified reporter molecules may unintentionally affect biological analysis through unforeseen responses. Moreover, the use of reporter molecules further increases both the complexity and cost of the overall system.

Image analysis-based microfluidic screening has emerged as a powerful approach for high-throughput screening (HTS), particularly in biological assays involving cell deformation^10^. Several deformation-based approaches have been employed to assess cell deformability under various conditions (mechanotyping), revealing consistent cellular responses to osmotic stress across platforms. These studies also highlighted method-specific sensitivities to deformability changes resulting from cytoskeleton disassembly. Additionally, the recent introduction of real-time monitoring systems^11^ and the application of deep learning-based image sorting^12,13^ offer promising advancements for future development of ultra-high throughput screening (uHTS) methods. However, despite this progress, image-based, label-free protocols to monitor the progress of biochemical reactions inside droplets have not yet been demonstrated.

While active systems, such as FADS, remain widely applied, recent studies introduced passive microfluidic methods that enable droplet screening based on intrinsic physical properties^14–17^.These protocols eliminate the need for fluorescent probes, dyes, and complex optical or electronic equipment. The transition toward passive systems enhances accessibility and cost-efficiency while retaining the ability to perform high-throughput and single-cell analyses. These advancements represent a significant step forward in the development of user-friendly and scalable microfluidic screening platforms^18^. However, in contrast with active systems for droplet sorting, passive methods often lack real-time monitoring capabilities and do not generate raw data for subsequent downstream analysis, thereby limiting their broader applicability. A notable example is the recently developed deformability-based passive droplet sorter (DPDS), designed for the label-free screening of microbial proteolytic activity^19^. Although the DPDS demonstrated effectiveness in isolating microbial strains through a barrier-based sorting mechanism, this device did not enable the quantification of proteolytic activity.

To overcome the limitations of both image-based and passive droplet screening systems, we have developed a method that employs image analysis to quantitatively monitor changes in the physicochemical properties of droplets. We then demonstrated how this label-free approach can be used for the ultra-high-throughput evaluation of biocatalytic processes within picoliter droplets. As a model, we investigated the growth of proteolytic microbial consortia in gelatin-laden droplets. The digital imaging protocol enables real-time detection and quantification of droplet deformation, which directly correlates with the extent of protein substrate degradation inside the droplets. This automated and robust platform offers a high-throughput, label-free solution for microbial protease discovery and application in industrial and environmental contexts. The research focused on four core objectives: i. designing a microfluidic to induce droplet deformation, ii. developing an automated workflow for individual droplet detection and shape analysis, comparing standard image analysis and AI-based methods, iii. validating the newly developed image-based system, and iv. screening of an environmental sample.

This microfluidic assay integrates high throughput, quantitative precision, and ease of use. Remarkably, the method achieved superior performance through single-cell encapsulation and screening, yielding higher colony-forming units (CFU/mL) of both proteolytic and non-proteolytic strains compared to traditional cultivation on solid selective media.

## Materials and Methods

### Fabrication of microfluidic devices

The microfluidic devices employed in this study were designed using AutoCAD (Autodesk), and master molds were fabricated using standard photolithography techniques. Polydimethylsiloxane (PDMS) devices were subsequently produced via soft lithography. A detailed description of the fabrication procedure is provided in the Supporting Information.

### Gelatine droplet generation and cultivation of microbes

Bacterial cells (OD ∼0.5) from an overnight shaking in physiologic solution were resuspended in rich-gelatine medium to the desired concentration. Microemulsions were generated using a flow-focusing droplet generator with an oil phase containing 5% fluorosurfactant at 25°C (RAN). The emulsions were collected in a droplet chamber and incubated at 40°C. A peristaltic pump supplied oxygen dissolved in the oil to the bacteria inside the droplets^20^, while preventing air bubbles from entering. Complete protocols for droplet generation and microbial cultivation are available in the Supporting Information.

### Droplet generation and bead labelling

To generate droplets with different gelatine concentrations, each identifiable during video analysis, we labelled each concentration with varying amounts of polystyrene beads (∼1 µm diameter). First, the stock solution (LB 0.5× + 7.5% gelatine) was supplemented with 0.5% (v/v) beads. Lower gelatine concentrations were subsequently prepared by diluting this stock solution with LB 0.5×. Droplet generation conditions were identical to those described above; detailed protocols are provided in the Supporting Information.

### Isolation of microbial samples

To prepare wastewater sludge samples for further experiments, aliquots of sludge were collected from containers stored in the cold room and transferred into 50 mL centrifuge tubes. Samples were centrifuged at 2000 rpm for 2 minutes to precipitate the particulate matter. After centrifugation, 1 mL of the supernatant was resuspended in 100 mL of physiological saline solution in a 0.5 L sterile Erlenmeyer flask. The flask was incubated at 25°C with continuous shaking at 250 rpm for 24 hours to ensure thorough interaction of sludge components with the solution. The suspension was then filtered through a 40 µm mesh filter to remove larger particles, and the filtrate was collected for subsequent cell encapsulation and screening on solid selective medium (skimmed milk agar).

### Bacteria cultivation media

The skimmed milk medium used for bulk screening consists of the following components: agar 15 (g/L), yeast extract 1, peptone 2, and skimmed milk powder 30. The droplet cultivation medium was prepared by mixing preheated gelatine (75 g/L final concentration), with LB medium (0.5× final concentration). Bacteria were gently mixed by pipetting the gelatine medium to minimize foam formation.

### Image analysis

The IDO analysis is based on an optimised Python script that processes high-speed video footage to extract and analyse shape-based features of moving particles within a defined region of interest (ROI). The script performs background subtraction, contour detection, and morphometric analysis to obtain shape descriptors for each detected droplet, such as area, perimeter, and AR. It also saves annotated video frames and quantitative data. Finally, it clusters data from temporally adjacent frames to compute average shape descriptors and their standard deviations, which are exported for further analysis.

Traditional approaches to image processing, such as intensity thresholding or edge detection, often rely on manually selected thresholds and filters. These limitations become particularly significant in high-throughput analyses, where not only accuracy but also scalability and automation of the process are required. In this work, a convolutional neural network (CNN) model based on one of the most efficient architectures - YOLO (You Only Look Once) - was used for image analysis. The YOLO architecture differs from traditional approaches in that it transforms the detection problem into a single regression task, without the need to generate region proposals. The input image is divided into a grid, and for each droplet, bounding boxes and class probabilities are predicted. YOLO processes the image in a single pass, providing very high processing speed while maintaining high accuracy, even in cases with multiple overlapping objects of potentially different shapes^21^. Before applying machine learning, a script was written in Python 3 (all scripts can be found on GitHub: https://github.com/Microfluidic-UW/Induced_Droplet_Ovalisation) to detect a moving droplet in a selected region of interest (ROI) in the video, apply a binary mask to the droplet, and save the video frame. Droplet motion was detected using background subtraction (MOG2), followed by Gaussian blur and binary thresholding to enhance the segmentation. Contours were then extracted and filtered based on area, circularity, and aspect ratio (AR) to isolate droplet-like shapes. For each valid detection, a red mask was applied to the original frame, and the corresponding bounding box coordinates were calculated. These values, along with the droplet’s class label, were saved in a YOLO-compatible .txt file format for training the YOLOv11s model.

## Results and Discussion

Proteolytic microorganisms represent a key class of strains in biotechnology, playing an important role in industries such as bioenergy, food processing, and pharmaceuticals^22–25^. Traditional methods for screening proteolytic activity, such as halo-based assays on agar plates like skimmed milk agar^26,27^, rely on the formation of clear halos around proteolytic colonies. However, these techniques are limited by low throughput, significantly reduced growth of microbial strains on solid media^28^, and susceptibility to false positives^29^. Additionally, the reliance on subjective interpretation of halos further undermines their efficiency, making this approach only partially effective for screening of microbial communities. Microfluidic screening of proteolytic activity has been demonstrated in the literature, but it has been predominantly focused on mammalian cells, employing fluorescence-based assays^30–32^.

A commonly used screening method employs a gelatin-rich medium that serves both as a protein substrate and a solidifying agent for detecting various classes of proteases, including metalloproteases, serine, aspartic, and cysteine proteases. Proteolytic activity is typically indicated by the liquefaction of the medium, resulting from enzymatic degradation of gelatin into soluble fragments^19^. We introduce a microfluidic-based method for the selective screening of proteolytic microbes, consisting of three key steps. First, individual microbial cells are encapsulated in picoliter-scale gelatin droplets using a flow-focusing microfluidic device. Following incubation, the droplets are reinjected into the IDO device, where a high-speed camera captures droplet deformation in a high-throughput, automated manner, as illustrated in Figure 1.

**Figure 1.**
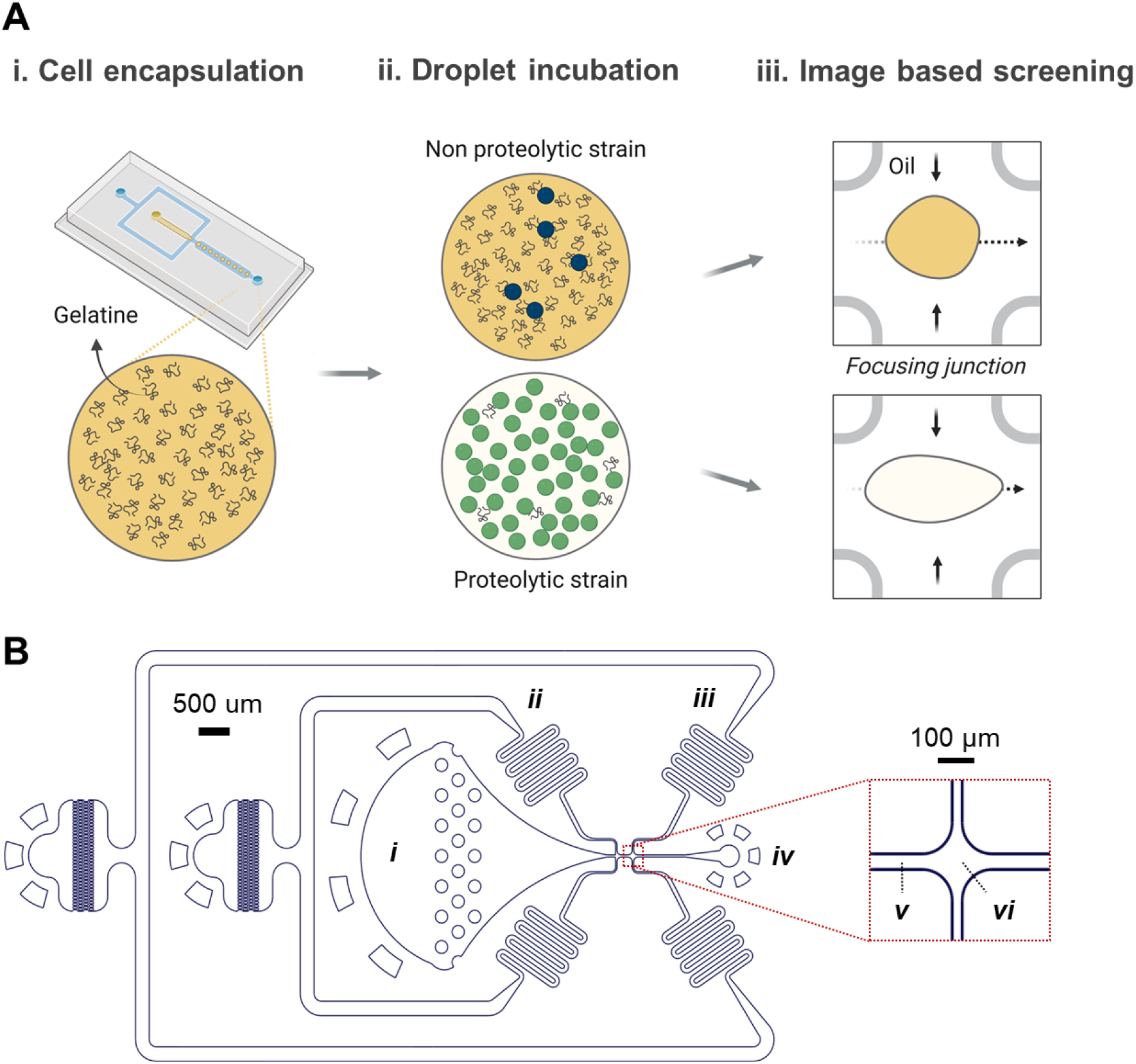
Detection of microbial proteolytic activity via droplet deformability image-based assay. This schematic illustrates a high-throughput, label-free method for screening proteolytic strains based on droplet viscoelasticity changes. **Panel A:** The platform enables the screening of microorganisms based on enzymatic activity, offering a powerful tool for biotechnological applications. First, a flow-focusing microfluidic device encapsulates single bacterial cells in picolitre-sized gelatine droplets. Non-proteolytic strains (top) maintain the integrity of the gelatine matrix, preserving the original viscoelastic properties of droplets. In contrast, proteolytic strains (bottom) hydrolyze gelatine during clonal growth and liquefying the droplets. Droplet deformability is assessed via the IDO device at a flow-focusing junction. Non-proteolytic droplets remain spherical under the influence of squeezing oil (black arrows), while proteolytic droplets become elongated. **Panel B:** The architecture of the device is displayed in the blue schematics. The design of the chip consists of the following key components: i.reinjection chamber, ii. spacing oil channel, iii. channel for squeezing oil, which induces droplet deformation, iv. device outlet, v. main channel where droplets flow leading from the reinjection chamber, vi. region of interest (ROI), where droplet imaging and analysis take place

### The design of the deformability-based device

The single-layer, 50 μm-deep IDO device used in this study - shown in Figure 1 - represents an advancement over the previously developed double-layer DPDS chip. Droplets, following incubation in an external tubing system, first enter the reinjection chamber of the device. This chamber tapers into a 50 μm-wide main channel and connects to a reinjection module incorporating a double flow-focusing junction that introduces two sequential oil streams. The first junction uniformly spaces and accelerates the droplets, directing them toward a geometrically symmetrical squeezing junction where real-time image analysis is conducted. Positive droplets (low gelatin content) are easily deformed in the junction, whereas negative droplets (high gelatine content) remain solid, passing through the squeezing junction unmodified and retaining their spherical shape.

### High-throughput generation and dynamic incubation of droplets for a deformability-based assay

As demonstrated in Video S1, an aqueous solution containing 7.5% gelatine is first emulsified into a monodisperse droplet suspension (∼1.5-2 kHz, 100 pL) at 25 °C using a flow-focusing junction. The droplets containing bacteria are then incubated at 40 °C for 36 hours under dynamic conditions (0.6 mL/h, with 5% fluorosurfactant in HFE-7500)^20,33^. For high-throughput screening (HTS) using the IDO device, a temperature range of 20–25 °C is required. Our observations indicate that conducting the assay below 20 °C typically results in liquid droplets being fragmented by solid ones during reinjection. In contrast, temperatures exceeding 25 °C result in uniform deformation of all droplets, regardless of gelatin content, thereby eliminating the assay’s ability to distinguish between proteolytic and non-proteolytic samples under the optimized conditions.

### Automated droplet detection

The optimized Python script automates video-based image analysis to quantify droplet deformability. The workflow involves video processing, morphological feature extraction, and statistical analysis of the collected data. The script first loads a video file, extracts metadata such as the total number of frames and frames per second, and applies a region of interest (ROI) to isolate relevant areas of the microfluidic channel. A background subtraction model (MOG2) enhances contrast, followed by Gaussian filtering and thresholding to generate a binary mask. Particle contours are identified, and only those with a set area interval are considered for analysis, i.e. 8500 - 10500 pixels. This range excludes smaller objects such as satellite droplets or debris, as well as larger ones as merged droplets. Morphological features are computed, including area, perimeter, bounding box dimensions, AR, circularity, convexity, elongation, solidity, rectangularity, extent, and Feret diameter, which is estimated as the maximum object length based on the relationship between area and perimeter. The extracted parameters (raw data) are stored in a Pandas DataFrame and exported as an Excel file (particle_data.xlsx) for further processing. The recorded particle data is subsequently grouped based on frame number using a clustering approach, and statistical evaluations are performed by calculating mean values and standard deviations within a specified frame range. The processed dataset is saved as an additional output file (processed_data.xlsx), providing a comprehensive characterization of droplet deformation as shown in Figure 2, additional information on droplet shape descriptors we calculated can be examined in Figure S2.

**Figure 2.**
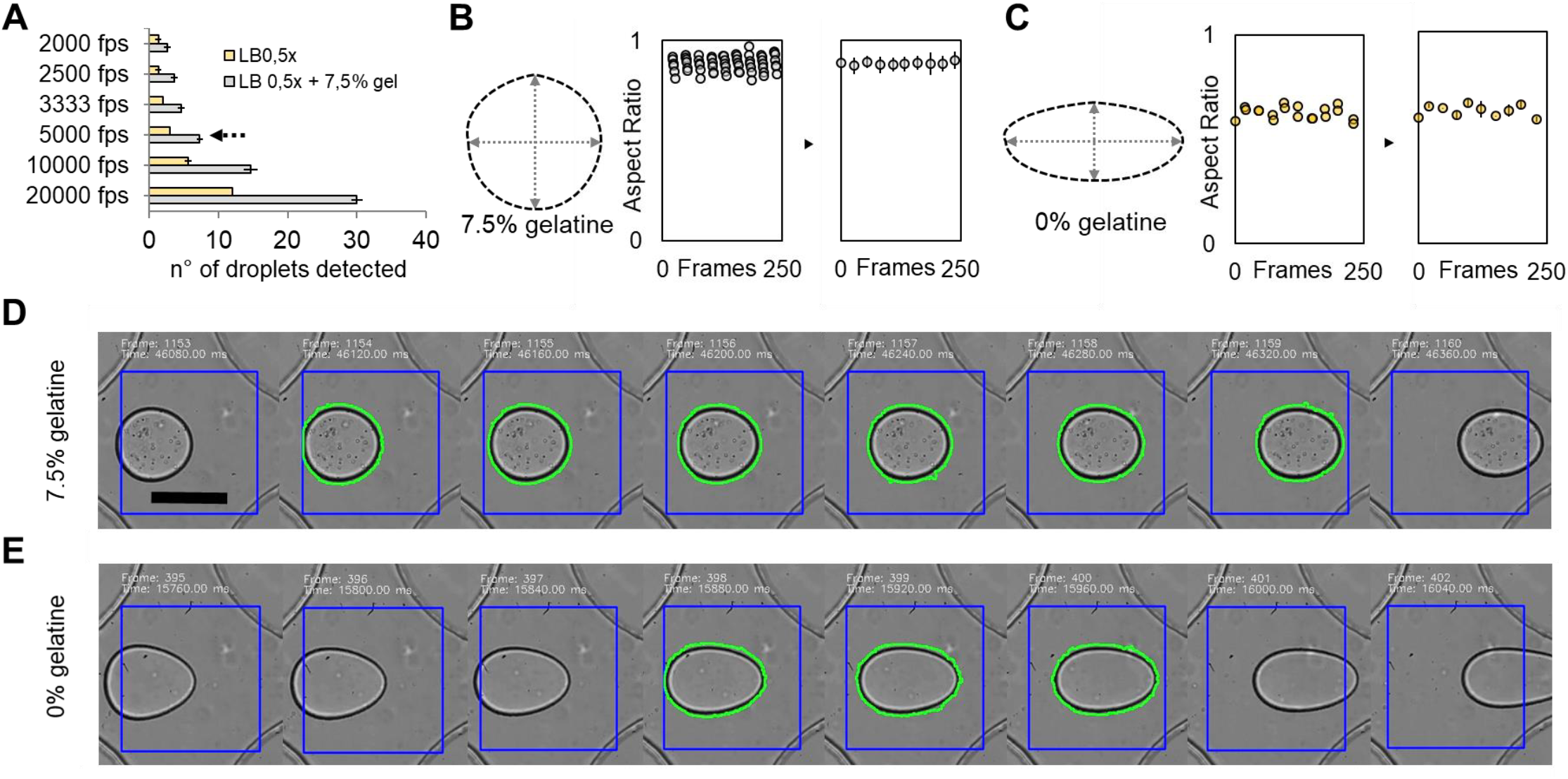
Automated detection of droplets and data processing. This figure illustrates the image-based workflow used to assess droplet deformability and optimize high-speed imaging parameters. **Panel A** shows a bar plot evaluating the effect of frame rate (frames per second, fps) on droplet detection efficiency. Grey bars represent negative droplets (LB 0.5× with 7.5% gelatin), while yellow bars correspond to positive droplets (LB 0.5× without gelatin). The black arrow indicates the frame rate selected for downstream analysis. **Panels B-C:** By combining this frame rate with the proposed screening throughput, we ensured the detection of multiple frames for both negative (**B**) and positive droplets (**C**). The difference in the number of frames detected between negative and positive droplets is attributed to their distinct deformability, which leads to varying residence times as they transit through the region of interest (ROI). To differentiate the two droplet populations, negative droplets were labeled with 0.5% (v/v) of ∼1 µm diameter polystyrene beads. Only droplets fully contained within the region of interest were detected (ROI, blue square; scale bar 50 µm), as highlighted by the green circle overlaying the droplet perimeter. **Panels D-E:** The bottom part of the figure presents a schematic representation of negative (**D**) and positive droplets (**E**) as they pass through the ROI. Their final AR values were calculated as the average across detected frames.

### Influence of oil flow rates and gelatine concentrations on droplet deformability

The proposed assay relies on droplet deformation as a function of gelatine content. Droplets with low gelatine content (positive) are liquid, whereas those with higher gelatine content (negative) are solid. On-chip deformation is induced by the flow of oil introduced in the second junction from the side channels of the main microfluidic channel. The extent of deformation varies significantly across the 0–7.5% gelatin concentration range and is strongly influenced by the squeezing oil flow rate. To evaluate the impact of flow rates on droplet deformability, we analyzed shape descriptors such as the AR across a range of squeezing oil flow rates (0.3-1.8 mL/h), as shown in Figure 3. Our results indicate that for 0-7.5% gelatine droplets, an optimal flow rate of 1.2 mL/h maximizes the range of AR, with a minimum of 0.5-0.6 and a maximum of 0.9-1.0. At lower flow rates, deformation is insufficient for meaningful AR analysis, whereas at higher flow rates, both negative and positive droplets deform similarly, diminishing the assay’s sensitivity to differences in gelatin content.

**Figure 3.**
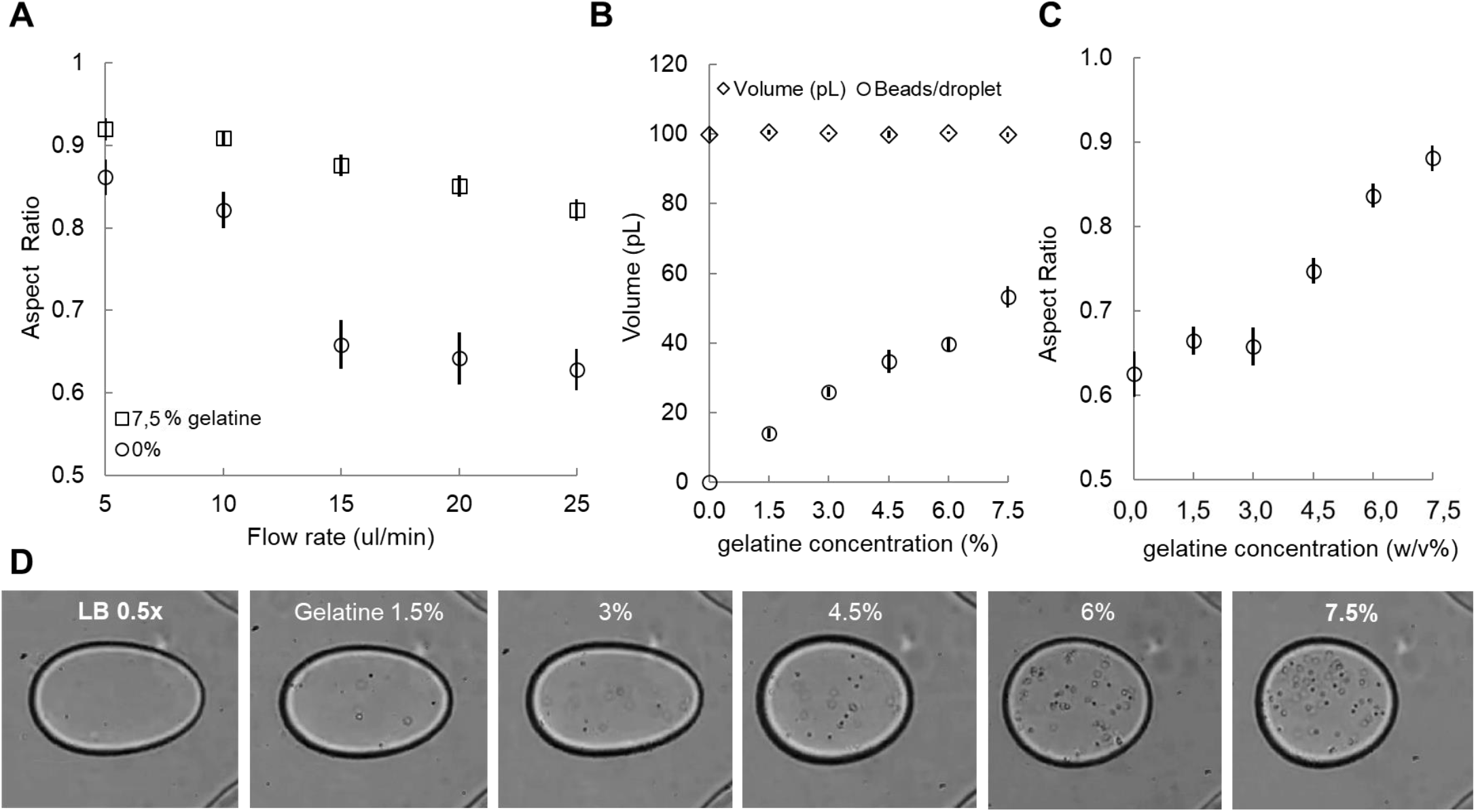
Influence of oil flow rate and gelatine concentration on droplet AR. This figure demonstrates how oil flow rate and gelatin concentration influence droplet deformability, as measured by AR. A decrease in AR reflects increased deformation and a more elongated droplet shape under shear stress from the squeezing oil flow. **Panel A:** Initially, we tested two different compositions 7.5% gelatine (squares) and 0% gelatine (circles), across a range of flow rates (5-25 µL/min) to determine the optimal flow rate that maximises the measurable difference between negative and positive droplets. The greatest separation in AR was observed at 20 µL/min, identified as the optimal flow rate for the assay. **Panel B**: We proceeded to test a series of droplets with different gelatine concentrations. Each concentration was labelled with a different amount of polystyrene microbeads, as shown in the scatter plot. The diamonds represent droplet volume (pL), which remained stable at approximately 100 pL across different gelatine and beads concentrations. The circles indicate the number of beads per droplet, showing a decreasing trend as gelatine concentration is reduced. **Panel C:** Higher gelatine concentrations resulted in reduced droplet deformation, leading to a high AR, while lower gelatine concentrations exhibit greater deformation, reaching a minimum AR of approximately 0.6. **Panel D**: Micrographs of droplets with varying gelatine concentrations are shown in the bottom part, ranging from LB 0.5× to 7.5% gelatine droplets. These images illustrate increased droplet stiffness and reduced deformation with increasing gelatine concentration.

To further investigate the relationship between gelatin content and droplet deformability, we conducted a screening of multiple LB 0.5x droplets with varying gelatine concentrations. To accurately detect each concentration, we labelled the 7.5% gelatine medium with 0.5% (v/v) of ∼1 µm diameter polystyrene microbeads (Sigma-Aldrich) - lower gelatine concentrations were then prepared by serial dilution. The generation of 100 pL droplets was not affected by the presence of microbeads, and the number of beads per droplet was estimated by counting them in a set of droplets from video recordings for each concentration. Screening droplets with different gelatine concentrations demonstrated a linear relationship between gelatine content and deformation within the 7.5-3% range. However, droplets with gelatin concentrations below 3% exhibited similar deformation behavior under fluid dynamic stress at the IDO device junction, indicating a loss of discriminatory power at lower concentrations.

To develop an automated classification system for droplet deformability, we trained a YOLO model using six videos from microfluidic experiments involving droplets with known gelatin concentrations. Droplets with 7.5% gelatin were spherical in shape and served as analogs for either empty droplets or those containing non-proteolytic bacteria. In contrast, droplets with gelatin concentrations approaching 0% simulated those containing proteolytic bacteria capable of degrading the gelatin matrix. From these videos, 2,550 frames were extracted and annotated with overlaid masks and corresponding .txt files, which were used to generate the training dataset. The dataset was randomly split, with 80% (2,040 images) allocated for training and 20% (510 images) reserved for validation after each epoch. The model was trained over 100 epochs on a Google Colab virtual machine equipped with a T4 GPU, with a total training time of approximately two hours. Next, the model was validated using a separate set of six videos containing droplets with varying gelatine concentrations: 0, 1.5, 3, 4.5, 6, and 7.5%. For videos with 0% and 1.5% gelatin concentrations, all detected droplets were classified as potentially proteolytic. For droplets at a 3% concentration, 97% of droplets were identified as proteolytic, while 3% were classified as non-proteolytic. In contrast, videos containing droplets with 4.5%, 6%, and 7.5% gelatin were consistently classified as non-proteolytic. These results are consistent with microfluidic observations: droplets with 0–3% gelatin exhibit similar deformation behavior under fluid dynamic stress, while differences in shape and size become less distinguishable at higher gelatin concentrations.

### Screening of a proteolytic strain from a synthetic microbial community

To evaluate the proposed microfluidic method under microbial culture conditions, we encapsulated single cells in picolitre droplets and cultivated them using two reference strains: a highly proteolytic *Pseudomonas aeruginosa* isolated from wastewater sludge and *Escherichia coli* as a non-proteolytic control strain. The emulsion primarily consisted of empty droplets (∼80%) to minimize the probability of multiple cells being encapsulated within the same droplet. Each strain was cultured separately overnight, diluted, and then mixed to simulate a synthetic environmental microbial community. The cell mixture was suspended in LB 0.5× supplemented with 7.5% gelatine to achieve a final concentration in droplets of approximately λ = 0.1 per strain. The droplets were incubated off-chip, and every 12 hours, microcultures were injected into the IDO device to assess microbial proteolytic activity, Figure 4. *P. aeruginosa* exhibited significantly faster growth in droplets compared to the non-proteolytic strain. While *E. coli* showed minimal growth, its distinct colony morphology enabled clear classification of all droplet types: i. empty droplets, ii. *E. coli* droplets, iii. *P. aeruginosa* droplets.

**Figure 4.**
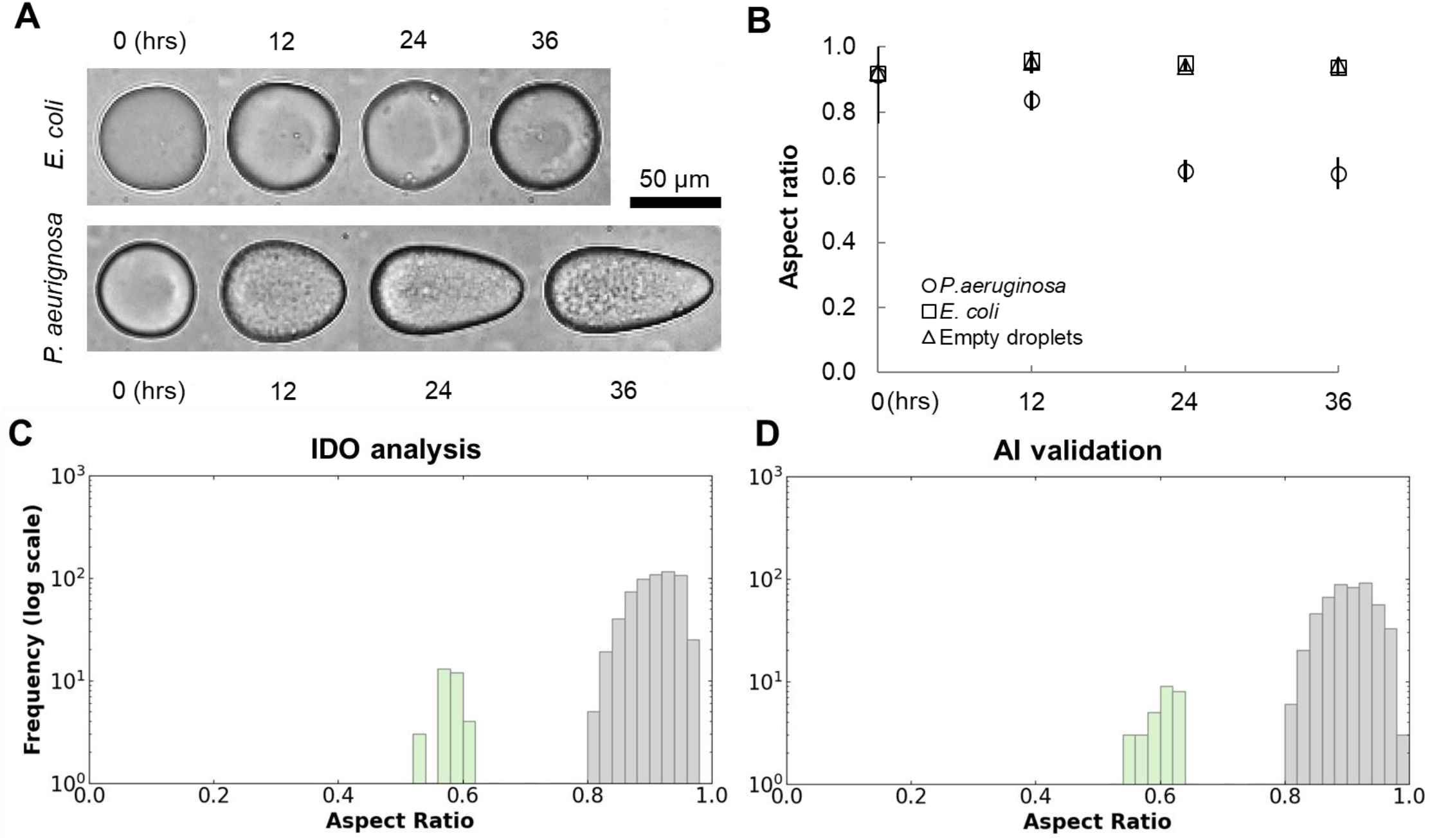
IDO method for screening microbial proteolytic activity. Clonal cultivation of strains originating from single cells encapsulated in LB 0.5× supplemented with 7.5% gelatine droplets was monitored in a time-course experiment every 12 hours. **Panel A:** Micrographs of droplet cultures illustrate changes in droplet shape and culture morphology of reference strains during image-based screening. Growth rates varied significantly, with *P. aeruginosa* being the only bacterium capable of metabolising gelatine. **Panel B**: The proteolytic activity of the two-strain community was successfully detected starting from the 12^th^ hour of incubation. **Panel C:** The histogram on the left shows the distribution of *P. aeruginosa* (λ = 0.1), based on droplet AR measured by IDO analysis. Green bins represent values below 0.65 after 24 hours of incubation. **Panel D:** Similarly, the AI analysis of droplets confirmed the image-based quantification of proteolytic *P. aeruginosa* present in the mock community.

We next analyzed the droplets using two complementary approaches: standard image analysis to calculate AR and a machine learning-based classification model. Automated image analysis using a Python-based script detected 630 droplets, correctly identifying all 33 proteolytic-containing droplets with aspect ratio (AR) values below 0.65, and no false positives. This corresponds to a positive fraction of P. aeruginosa of 5.23%. The machine learning model performed 528 detections in total, of which 33 droplets were classified as containing proteolytic bacteria-corresponding to approximately 6.25% of the total. This result aligns with expectations based on the experimental loading conditions. The model’s confidence in droplet classification ranged from 70% to 94%, with the majority of detections falling within the upper end of this range, indicating reliable performance across the analyzed dataset. For each droplet detected by the machine learning (ML) model, we also quantified its AR and compared these values to those obtained through standard automated image analysis. The results are presented side by side in **Figure 4C–D**, highlighting the consistency between both approaches.

### Environmental screening of wastewater sludge

The sludge sample we studied was collected from a wastewater treatment plant in Wołomin, Poland, where biogas is produced using sewage sludge as a sustainable energy source. To maintain optimal conditions for microbial activity, the substrate is recirculated at 40 °C in the biogas facility. Our novel microfluidic protocol was first applied to monitor proteolytic bacteria in wastewater sludge. Proteolytic bacteria facilitate substrate solubilization by breaking down complex proteins into bioavailable peptides and amino acids, thereby enhancing subsequent acidogenesis and methanogenesis. This step is crucial for optimizing the hydrolysis phase, the rate-limiting step in anaerobic digestion. The sludge was first resuspended as described in the Materials and Methods section, followed by single-cell encapsulation in 100 pL droplets (λ = 0.2). In parallel, conventional screening methods were employed, including use of LB and skimmed milk agar medium, as shown in Figure 5. Image analysis was performed at a throughput of 0.5 kHz within the ROI. The syringe pump settings were set as follows: sludge sample - 0.12 mL/hour, spacing oil containing 2% fluorosurfactant - 0.3 mL/hour, squeezing oil, without surfactant - 1.2 mL/hour.

**Figure 5.**
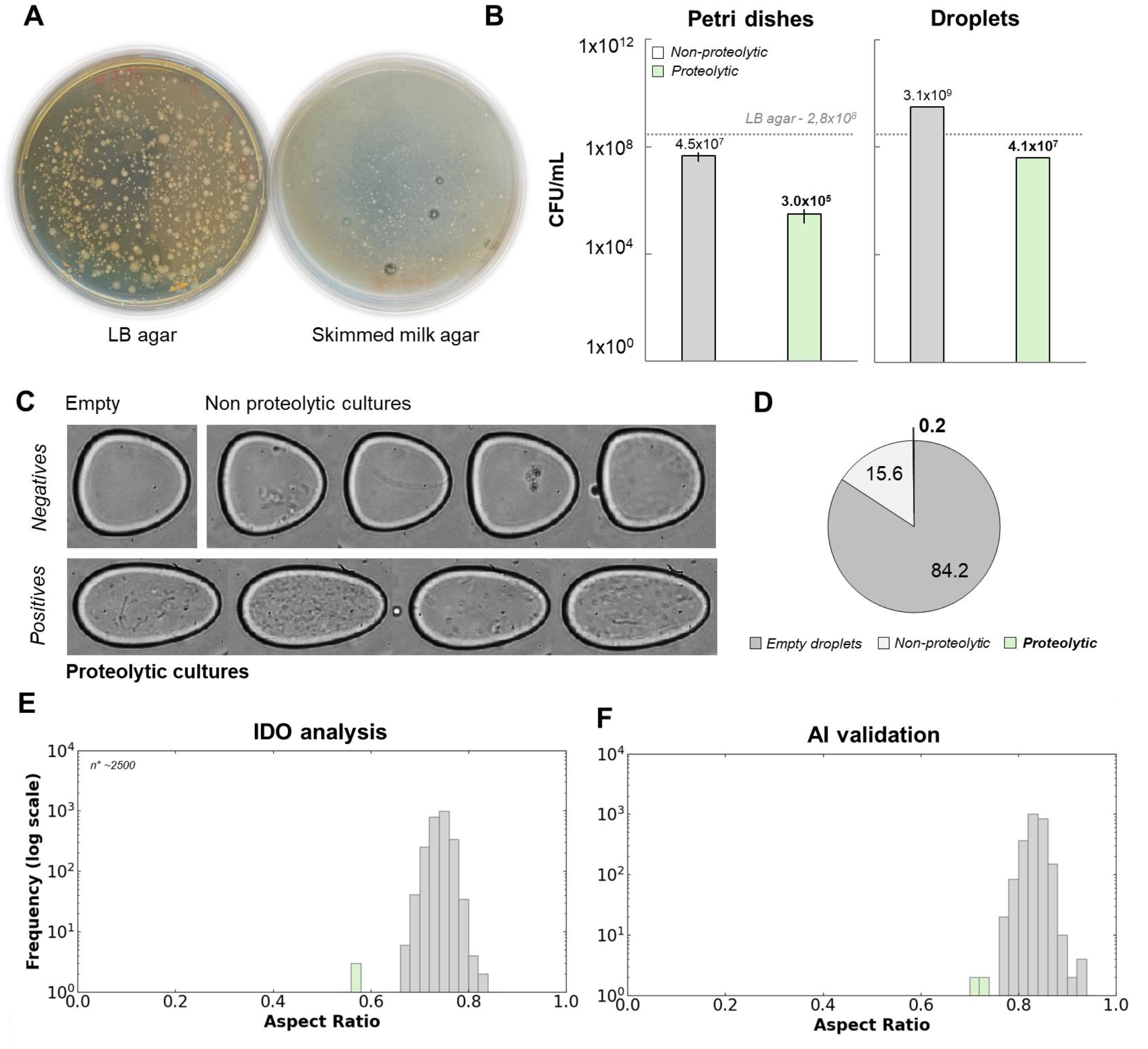
IDO-HTS of proteolytic consortia in wastewater sludge. **Panel A:** The top right corner presents photographs of traditional bacterial screening methods, including LB agar for cultivating heterotrophic microbes and skimmed milk agar for selectively enumerating proteolytic colonies. Screening was performed in triplicate for three dilutions. CFU concentration in wastewater sludge was assessed upon cultivation on solid media. **Panel B:** Bar charts, reporting figures from both Petri dish and droplet-based screening, show a significant increase in CFU detection compared to traditional solid media protocols. This highlights the superior screening performance of our method for cultivating both non-proteolytic and proteolytic strains. **Panel C**: Droplet-based screening highlights the different morphology of empty droplets, non-proteolytic colonies (negatives), and proteolytic colonies (positives) originating from single-cell encapsulation. **Panel D**: The pie chart illustrates the droplet emulsion composition: empty droplets (dark grey), non-proteolytic colonies (grey), and proteolytic colonies (green). Most droplet cultures (15.6%) contained non-proteolytic strains, while 0.2% exhibited proteolytic activity, non-proteolytic cultures encapsulated in droplets were quantified manually from videos. Proteolytic activity was detected through significant variations in AR, indicating changes in droplet morphology. **Panels E-F**: The left histogram shows the AR distribution of droplets identified by IDO analysis (**E**), while the right histogram confirms the environmental screening results through AI analysis (**F**).

The microbial proteolytic activity of this consortium was screened after 24 hours of incubation. Approximately 2500 droplets were analyzed, requiring a volume of 2.5 × 10^−7^ mL—significantly lower than the 15 mL required for a single 90 mm Petri dish in conventional assays. For standard plating, three replicates at three serial dilutions were performed for both cultivation media, resulting in an estimated total volume of ∼270 mL. IDO screening revealed a significant increase in the number of strains cultured in droplets, both proteolytic and non-proteolytic bacteria. This highlights the advantages of single-cell encapsulation in enhancing the sensitivity and throughput of environmental microbial screening compared to traditional cultivation methods.

Automated image analysis using a Python-based script detected 2453 droplets, correctly identifying all 4 proteolytic-containing droplets with no false positives. The same video data was analyzed using a trained classification model. The model processed 2,499 droplets and identified 4 as containing proteolytic bacteria. Classification confidence ranged from approximately 70% to 96%, indicating the model’s level of certainty in detecting gelatin-degrading activity within individual droplets.

## Conclusion

Despite the challenges in developing novel accurate tools for screening environmental samples, droplet microfluidic techniques have already demonstrated significant applications in modern microbiology and biotechnology^32^. In this study, we present a microfluidic protocol based on single-cell encapsulation and clonal cultivation in picoliter droplets to detect microbial proteolytic activity in complex and unknown samples, such as wastewater sludge. Detection is achieved through an automated, user-friendly, and customizable Python script that analyses multiple shape descriptors and provides images of droplets. The screening of proteolytic microbes is carried out using a novel chip device that employs high flows of continuous phase to induce gelatine droplet deformation. We optimized multiple conditions, enabling the analysis of up to 200 droplets per second. To enhance droplet detection efficiency, we performed a series of experiments to refine key parameters, including the influence of frame rate (fps) on single-droplet detection and the identification of the most informative shape descriptors for assessing microbial proteolytic activity. To support and validate the detection of proteolytic-positive droplets, we implemented an AI-based image analysis workflow. Deep learning models are capable of identifying subtle morphological changes in droplets, thereby reducing the need for manual inspection and enhancing both reproducibility and scalability of the screening assay^34^. Despite its advantages, deep learning comes with several challenges. It requires large, well-annotated datasets, significant computational power, and its decisions are often difficult to interpret. In contrast, conventional image analysis techniques, which rely on manual thresholding, are more accessible, computationally efficient, and easier to modify. However, they may struggle in scenarios involving complex or noisy backgrounds, where fine discrimination is required. Ultimately, the choice between AI-based and conventional approaches depends on the specific trade-offs among accuracy, accessibility, and operational feasibility in a given application. During the final phase of optimization, we evaluated various gelatin concentrations, screened a mock microbial community, and ultimately applied the droplet-based screening platform to an environmental sample. The results were benchmarked against those obtained through conventional isolation using solid cultivation media.

While droplet-based platforms such as AADS and FADS offer high throughput and sensitivity, they typically rely on fluorescent dyes or labeled substrates. In contrast, image-based droplet sorting presents a promising label-free alternative. Here, we introduce for the first time a high-throughput method for the qualitative monitoring of proteolytic bacterial growth and activity, based on changes in the physicochemical properties of droplets. Unlike droplet methods based on fluorescence assay, the proposed enrichment screening platform eliminates the need for costly or non-commercially available fluorogenic substrates to detect protein degradation^31^. Its simplicity makes it more user-friendly than classical methods like Petri dish screenings, requiring only oil flow adjustments for precise droplet spacing and accurate video analysis. Moreover, in comparison to previously reported passive systems for deformability-based droplet screening^15–17,19^, our method achieves significantly higher throughput, offering both speed and scalability for environmental microbiology applications.

Deformation-based beads and droplet analysis might be beneficial in many biotechnological applications. Although cell deformation and imaging have been used in mechanotyping assays^10,35,36^ the use of such approaches for microbial community screening via droplet microfluidics remains limited. A key limitation of our gelatin-based droplet method lies in the change of temperature required for droplet solidification. Our findings indicate that successful image analysis depends on maintaining temperatures between 20–25 °C, as the deformability of the gelatin matrix is temperature dependent. To address this, future improvements could involve using methacrylated gelatin as a protease substrate, which may offer greater thermal stability.

Nonetheless, this highlights a key feature of our workflow - the proposed system can be adapted for virtually any hydrogel screening, enabling the monitoring of polymerization or degradation using various shape descriptors. When integrated with sorting electronics, including a droplet sorter, and a droplet deposition system, this method could serve as an easily implementable and a robust platform for screening and enriching microbial consortia. By leveraging real-time image analysis and automated sorting mechanisms, the platform could efficiently isolate droplets containing microbial consortia with desired traits, significantly enhancing the precision and scalability of environmental and biotechnological screenings. While these enhancements lie beyond the scope of the present study, they represent promising directions for future methodological development.

Overall, our cost-effective, label-free, and accurate system facilitates the monitoring of environmental samples and has demonstrated superior performance in cultivating strains compared to conventional methods. Induced Droplet Ovalisation (IDO) enhances the applicability of microfluidic screening, expanding its potential for both research and practical applications in microbiology.

## Supporting information

- Detailed protocols for photolithographic microfabrication of microfluidic device molds, a profilometer scan image of the IDO module, image-based analysis scripts, droplet shape descriptors, and microbial growth results of environmental strains inside droplets (PDF).
- High-viscosity gelatine droplet generation (LB 0.5x + 7.5% gelatine), Video S1 (MP4).
- LB 0.5x and LB 0.5 + 7.5% gel flowing in the ROI of IDO device, Video S2-S3.
- Screening of a binary mock consortia, Video S4 (MP4).
- Screening of environmental sample, Video S5 (MP4).
- IDO analysis of wastewater sludge, Video S6.
- Frames from IDO analysis of wastewater sludge.
- CAD files containing designs of microfluidic chips (ZIP).

## Glossary

uHTS / HTS: ultra High Throughput Screening / High Throughput Screening
AR: Aspect ratio
Gel: Gelatine
IDO: Induced Droplet Ovalisationa
CFU: Colony forming unit
LB: Lysogeny broth
fps: Frames per second

## Acknowledgements

This research was funded by the National Science Centre, Poland (grants SONATA 2021/43/D/ST4/03291 and SONATA BIS no. 2023/50/E/ST4/00545), project. IDUB small grants funded by the University of Warsaw also contributed to the execution of this research work (grant no. 01/14-01-00/2023). We also thank Lukasz Drewniak and Mikolaj Wolacewicz for helping coordinate the sampling trips and Lukasz Kozon for the fabrication of the master mould of a microfluidic chip. Part of Figure 1 was prepared with https://www.biorender.com/.

## References

1. Alain K, Querellou J. Cultivating the uncultured: limits, advances and future challenges. Extremophiles. 2009;13:583–594.

2. Watterson WJ, Tanyeri M, Watson AR, et al. Droplet-based high-throughput cultivation for accurate screening of antibiotic resistant gut microbes. Elife. 2020;9:e56998. doi:10.7554/eLife.56998

3. Mahler L, Niehs SP, Martin K, et al. Highly parallelized droplet cultivation and prioritization of antibiotic producers from natural microbial communities. Kana BD, Dhar N, Sclavi B, eds. eLife. 2021;10:e64774. doi:10.7554/eLife.64774

4. Najah M, Calbrix R, Mahendra-Wijaya IP, Beneyton T, Griffiths AD, Drevelle A. Droplet-Based Microfluidics Platform for Ultra-High-Throughput Bioprospecting of Cellulolytic Microorganisms. Chemistry & Biology. 2014;21(12):1722–1732. doi:10.1016/j.chembiol.2014.10.020

5. Baret JC, Miller OJ, Taly V, et al. Fluorescence-activated droplet sorting (FADS): efficient microfluidic cell sorting based on enzymatic activity. Lab on a Chip. 2009;9(13):1850–1858. doi:10.1039/b902504a

6. Postek W, Garstecki P. Droplet microfluidics for high-throughput analysis of antibiotic susceptibility in bacterial cells and populations. Accounts of Chemical Research. 2022;55(5):605–615.

7. Beneyton T, Thomas S, Griffiths AD, Nicaud JM, Drevelle A, Rossignol T. Droplet-based microfluidic high-throughput screening of heterologous enzymes secreted by the yeast Yarrowia lipolytica. Microbial Cell Factories. 2017;16(1):18. doi:10.1186/s12934-017-0629-5

8. Gielen F, Hours R, Emond S, Fischlechner M, Schell U, Hollfelder F. Ultrahigh-throughput–directed enzyme evolution by absorbance-activated droplet sorting (AADS). Proceedings of the National Academy of Sciences. 2016;113(47):E7383–E7389. doi:10.1073/pnas.1606927113

9. Medcalf EJ, Gantz M, Kaminski TS, Hollfelder F. Ultra-High-Throughput Absorbance-Activated Droplet Sorting for Enzyme Screening at Kilohertz Frequencies. Anal Chem. 2023;95(10):4597–4604. doi:10.1021/acs.analchem.2c04144

10. Urbanska M, Muñoz HE, Shaw Bagnall J, et al. A comparison of microfluidic methods for high-throughput cell deformability measurements. Nature Methods. 2020;17(6):587–593. doi:10.1038/s41592-020-0818-8

11. Neto JP, Mota A, Lopes G, et al. Open-source tool for real-time and automated analysis of droplet-based microfluidic. Lab Chip. 2023;23(14):3238–3244. doi:10.1039/D3LC00327B

12. Anagnostidis V, Sherlock B, Metz J, Mair P, Hollfelder F, Gielen F. Deep learning guided image-based droplet sorting for on-demand selection and analysis of single cells and 3D cell cultures. Lab Chip. 2020;20(5):889–900. doi:10.1039/D0LC00055H

13. Srikanth S, Dubey SK, Javed A, Goel S. Droplet based microfluidics integrated with machine learning. Sensors and Actuators A: Physical. 2021;332:113096. doi:10.1016/j.sna.2021.113096

14. Potenza L, Kozon L, Drewniak L, Kaminski TS. Passive Droplet Microfluidic Platform for High-Throughput Screening of Microbial Proteolytic Activity. Anal Chem. 2024;96(40):15931–15940. doi:10.1021/acs.analchem.4c02979

15. Staskiewicz K, Dabrowska-Zawada M, Kozon L, Olszewska Z, Drewniak L, Kaminski TS. Droplet microfluidic system for high throughput and passive selection of bacteria producing biosurfactants. Lab Chip. 2024;24(7):1947–1956. doi:10.1039/D3LC00656E

16. Pan CW, Horvath DG, Braza S, et al. Sorting by interfacial tension (SIFT): label-free selection of live cells based on single-cell metabolism. Lab Chip. 2019;19(8):1344–1351. doi:10.1039/C8LC01328D

17. Muta M, Kawakubo W, Yoon DH, et al. Deformability-Based Microfluidic Microdroplet Screening to Obtain Agarolytic Bacterial Cells. Analytical Chemistry. 2023;95(44):16107–16114.

18. Kaminski TS, Scheler O, Garstecki P. Droplet microfluidics for microbiology: techniques, applications and challenges. Lab Chip. 2016;16(12):2168–2187. doi:10.1039/C6LC00367B

19. Potenza L, Kozon L, Drewniak L, Kaminski TS. Passive Droplet Microfluidic Platform for High-Throughput Screening of Microbial Proteolytic Activity. Anal Chem. 2024;96(40):15931–15940. doi:10.1021/acs.analchem.4c02979

20. Mahler L, Tovar MA, Weber T, et al. Enhanced and homogeneous oxygen availability during incubation of microfluidic droplets. RSC Adv. 2015;5. doi:10.1039/C5RA20118G

21. Khanam R, Hussain M. Yolov11: An overview of the key architectural enhancements. arXiv preprint arXiv:241017725. Published online 2024.

22. Razzaq A, Shamsi S, Ali A, et al. Microbial proteases applications. Frontiers in bioengineering and biotechnology. 2019;7:110. doi:10.3389/fmicb.2023.1236368

23. Morellon-Sterling R, El-Siar H, Tavano OL, Berenguer-Murcia Á, Fernández-Lafuente R. Ficin: A protease extract with relevance in biotechnology and biocatalysis. International Journal of Biological Macromolecules. 2020;162:394–404. doi:10.1016/j.ijbiomac.2020.06.144

24. Niyonzima F, More S. Detergent-Compatible Proteases: Microbial Production, Properties, and Stain Removal Analysis. Preparative biochemistry & biotechnology. 2014;45. doi:10.1080/10826068.2014.907183

25. Craik CS, Page MJ, Madison EL. Proteases as therapeutics. Biochemical Journal. 2011;435(1):1–16. doi:10.1042%2FBJ20100965

26. Kasana RC, Salwan R, Yadav SK. Microbial proteases: detection, production, and genetic improvement. Critical reviews in microbiology. 2011;37(3):262–276. doi:10.3109/1040841X.2011.577029

27. Morris LS, Evans J, Marchesi JR. A robust plate assay for detection of extracellular microbial protease activity in metagenomic screens and pure cultures. Journal of microbiological methods. 2012;91(1):144–146. doi:10.1016/j.mimet.2012.08.006

28. Nosho K, Yasuhara K, Ikehata Y, et al. Isolation of colonization-defective Escherichia coli mutants reveals critical requirement for fatty acids in bacterial colony formation. Microbiology. 2018;164(9):1122–1132. doi:10.1099/mic.0.000673

29. Jones B, Sun F, Marchesi J. Using skimmed milk agar to functionally screen a gut metagenomic library for proteases. Letters in applied microbiology. 2007;45:418–420. doi:10.1111/j.1472-765X.2007.02202.x

30. Wu W, Zhang S, Zhang T, Mu Y. Immobilized Droplet Arrays in Thermosetting Oil for Dynamic Proteolytic Assays of Single Cells. ACS Appl Mater Interfaces. 2021;13(5):6081–6090. doi:10.1021/acsami.0c21696

31. Ng EX, Miller MA, Jing T, Chen CH. Single cell multiplexed assay for proteolytic activity using droplet microfluidics. Biosensors and Bioelectronics. 2016;81:408–414. doi:10.1016/j.bios.2016.03.002

32. Dhar M, Lam JN, Walser T, Dubinett SM, Rettig MB, Di Carlo D. Functional profiling of circulating tumor cells with an integrated vortex capture and single-cell protease activity assay. Proceedings of the National Academy of Sciences. 2018;115(40):9986–9991. doi:10.1073/pnas.1803884115

33. Neun S, Kaminski TS, Hollfelder F. Chapter Five - Single-cell activity screening in microfluidic droplets. In: Allbritton NL, Kovarik ML, eds. Methods in Enzymology. Vol 628. Academic Press; 2019:95–112. doi:10.1016/bs.mie.2019.07.009

34. Gardner K, Uddin MM, Tran L, Pham T, Vanapalli S, Li W. Deep learning detector for high precision monitoring of cell encapsulation statistics in microfluidic droplets. Lab on a Chip. 2022;22(21):4067–4080.

35. Chang Y, Chen X, Zhou Y, Wan J. Deformation-based droplet separation in microfluidics. Industrial & Engineering Chemistry Research. 2019;59(9):3916–3921.

36. An L, Ji F, Zhao E, Liu Y, Liu Y. Measuring cell deformation by microfluidics. Frontiers in Bioengineering and Biotechnology. 2023;11:1214544.

