## Supplementary Information file for "Induced Droplet Ovalisation (IDO): Image-based microfluidic method for high-throughput and label-free characterization of microbial proteolytic strains from wastewater sludge"

### Table of Content

#### Supplementary methods


| | | |
|---|---|---|
| i. | Fabrication of microfluidic devices | S1 |
| ii. | Photolithography | S1 |
| iii. | Soft lithography | S1 |
| iv. | Incubation chambers for droplets dynamic incubation | S2 |
| v. | Isolation of the proteolytic reference strain | S3 |
| vi. | Bacterial cultivation | S3 |
| vii. | Shape descriptors | S4 |


#### Supplementary figures


| | | |
|---|---|---|
| i. | Figure S1. Schematic of the dynamic incubation system | S1 |
| ii. | Figure S2. Quantitative analysis of droplet shape across different gelatine concentrations. | S4 |
| iii. | Figure S3. Validation and confidence in gelatine droplet detection. | |
| iv. | Figure S4. IDO analysis of wastewater sludge. | |


**Fabrication of microfluidic devices.** Every microfluidic devices used in this study were designed using AutoCAD (Autodesk), while the master molds were created through standard photolithography and soft lithography techniques[1]. The fabrication process of PDMS devices involved two main steps: first, the master molds with the designs were produced via photolithography; subsequently, PDMS was poured onto these molds to form replicas containing the microfluidic structures.

**Photolithography.** Microfluidic molds were fabricated on 3-inch silicon wafers (Microchemicals) using high-resolution acetate masks (Microlithography Services) and SU-8 series photoresist patterning (Kayaku Advanced Materials). The SU-8-coated wafers were subsequently exposed to UV light using an MJB4 mask aligner (SÜSS MicroTec)[2]. The CAD designs for the flow-focusing (FF) droplet generation device and the IDO (Induced Droplet Ovalisation) module are provided as a supplementary .dxf file. The structure thickness, which determines the final channel depth in the microfluidic devices, was measured using a GT-Contour profilometer (Bruker) with a 5x objective.

**Soft Lithography.** To fabricate a single microfluidic device, approximately 30 grams of PDMS (Sylgard 184, Dow) were measured into a plastic cup. The curing agent was then added at a 10:1 (w/w) ratio and thoroughly mixed before degassing in a vacuum chamber. The degassed PDMS was poured onto the SU-8 master wafer, placed in a Petri dish, and cured in an oven at 70°C for 4 hours. Once solidified, the PDMS layer was carefully peeled from the mold, and inlet and outlet holes were punched using a 1 mm biopsy puncher (Kai Medical). The flow-focusing droplet generators and IDO device chips were bonded to glass slides (VWR) using an automated plasma system (Zepto, Diener Electronics). The plasma treatment process involved three steps: first, a vacuum of 0.35 mbar was generated; next, oxygen was introduced into the chamber for one minute at a pressure of 0.5 mbar; finally, the plasma was activated at 30% power for 45 seconds. Hydrophobic modification of the chips was carried out by flushing the devices with a 0.5% solution of trichloro (1H,1H,2H,2H perfluorooctyl)-silane (Sigma-Aldrich) in Novec HFE-7500 oil (3M) and baked on a hot plate at 80°C for 30 minutes to evaporate fluorocarbon liquid.

**Incubation chambers for droplets dynamic incubation.** The storage chambers for droplet incubation were fabricated following the protocol described by Neun et al.[3]. A 1-mm biopsy punch (Kai Medical) was used to create perforations at the bottom tip and along the side of a 0.5 ml Eppendorf tube. The tube's lid was securely affixed to a 1-mm thick glass slide using cyanoacrylate glue (PR 1500, 3M). Subsequently, two 20-cm-long Teflon (PFTE) tubes (0.4 mm I.D., 0.9 mm O.D., Bola Bohlender) were inserted into the punched holes and adhered to the Eppendorf tube's surface. During droplet generation, the flow-focusing chip was connected to the upper tubing, facilitating the collection of droplets in the upper region of the tube due to their inherent buoyancy in such oil. Following incubation, to initiate droplet screening, the flow direction was reversed. oil was pumped through the side tubing, guiding the droplets toward the IDO reinjection chamber via the top tubing. The emulsions were incubated for 36 hours using dynamic droplet incubation (DDI), a method known to enhance microbial growth and metabolic activity[4]. The experimental setup employed in this study is illustrated in Fig. S2. Special care was taken to eliminate air bubbles from the tubing to ensure smooth operation. A schematic representation of the oxygenation system highlights the essential role of the peristaltic pump and bubble trap, which facilitated oxygen delivery while simultaneously preventing bubble formation within the incubation chamber. The pump operated with Tygon tubing (0.38 mm I.D., 0.90 mm O.D., Ismatec), which was connected and sealed to the incubation chamber using PTFE tubing (0.5 mm I.D., 1 mm O.D., Bola Bohlender). To maintain a stable, bubble-free environment, both ends of the tubing were submerged in a bubble trap - consisting of a 1.5 mL Eppendorf tube filled with 5% RAN fluorosurfactant in HFE-7500 - thereby completing the sealed system, Figure S1.

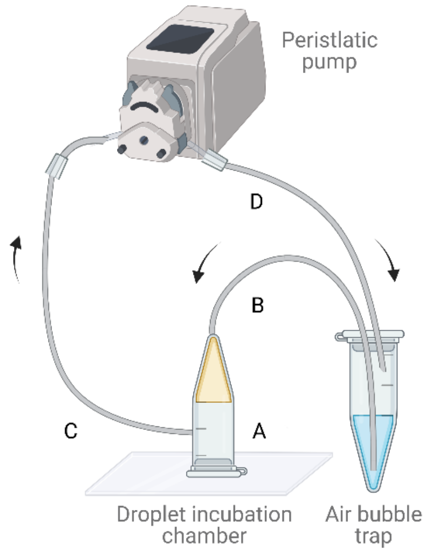


**Figure S1. Schematic of the dynamic incubation system.** The oxygenation system consisted of a closed-loop setup designed to maintain controlled oil flow. A 0.5 mL Eppendorf tube, serving as the droplet chamber, was affixed to a glass slide (A). A section of tubing was then attached to the upper part of the chamber (B), while a second tubing was inserted near the lid (C). Both tubing were securely connected using PTFE tubing and linked to a peristaltic pump (Reglo ICC, Ismatec) with adhesive glue. The other end of the tubing, directed from the pump, was attached to the lid of a second Eppendorf tube containing 5% fluoroSurfactant solution in filtered Novec HFE-7500 (D). This configuration ensured a continuous oil flow in a closed system. The oil was pumped at 0.6 mL/h, flowing from the top of the chamber - where droplets accumulated - toward the lower inlet.

**Isolation of the proteolytic reference strain.** The proteolytic reference strain used in this study was isolated from sludge collected at a wastewater treatment plant in Wołomin, Mazovia region, Poland. The sludge sample was resuspended in a physiological solution within 0.1 L flasks and incubated under shaking conditions (200 rpm) at 30°C for one hour. Following incubation, the sample was serially diluted and plated onto a skimmed milk medium to detect microbial proteolytic activity. Subsequent identification via 16S rRNA sequencing confirmed that the isolated strain was *Pseudomonas aeruginosa*.

**Bacterial cultivation.** For the development and validation of our microfluidic method, we employed two bacterial strains: i. *P. aeruginosa*, used to generate positive droplets containing a highly proteolytic bacterium, and ii. *E. coli* DH5α, a non-proteolytic strain serving as a negative control. Overnight cultures were prepared by inoculating a single colony of each strain into 10 mL of LB liquid medium in flasks, followed by incubation at 30°C with shaking at 100 rpm. Microbial growth was monitored by measuring the optical density at 600 nm (OD600) of 200 µL aliquots using a plate reader (Sunrise, Tecan) and Magellan™ software. The OD600 values obtained from the overnight LB cultures were used to calculate cell concentrations and adjust the inoculum for the desired cell-to-droplet occupancy. Bacterial cultures were collected during the exponential growth phase (OD600 ~0.5), ensuring that the majority of cells were viable and actively dividing, thereby improving the accuracy of cell encapsulation calculations. The droplet cultivation medium was prepared from two stock solutions: a gelatine stock (150 g/L gelatine) and an LB medium. The gelatine stock was preheated to 40°C to liquefy and diluted to achieve a final concentration of 75 g/L (7.5% gelatine). The final medium composition consisted of LB 0.5x supplemented with 7.5% gelatine. To minimize foam formation, bacterial cultures were first mixed with LB by vortexing before being gently combined with the gelatine solution through careful pipetting.

**Shape descriptors.** In addition to the aspect ratio (AR), which was used as a deformability descriptor for detecting microbial proteolytic activity, several other parameters were analyzed. To assess the monodispersity of the emulsion beyond visual inspection, droplet area and perimeter were measured. The results indicate that increasing gelatine concentration affects droplet shape and size. A notable linear correlation was observed between gelatine content and various shape descriptors, excluding AR, highlighting the role of gelatine in modulating droplet deformability, as shown in Figure S2.

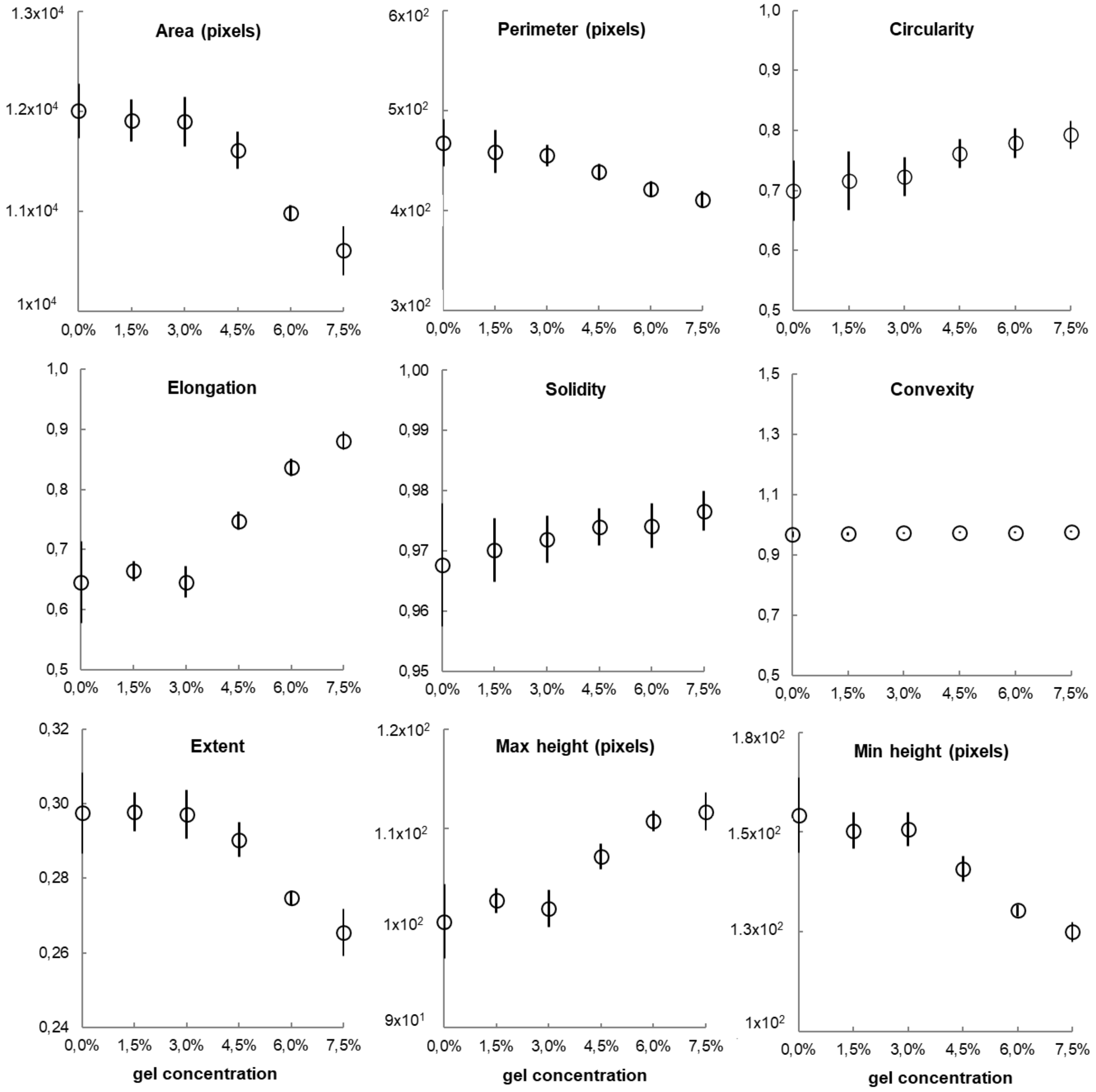


**Figure S2: Analysis of droplet shape across gelatine concentrations.** The figure presents a set of shape descriptors for droplets at varying gelatine concentrations, illustrating the influence of gelatine content on droplet morphology. The parameters analyzed included area, perimeter, circularity, convexity, elongation, solidity, extent, maximum height and minimum lenght. The area and perimeter values exhibit a trend correlating with increasing gelatine concentration, suggesting a change in droplet size. Circularity varies with gelatine content, especially at higher concentrations. Elongation increases, indicating greater deformation, while solidity decreases at lower gelatine concentrations and convexity remains stable. Extent decreases with higher gelatine content, while maximum height and minimum length depend on gelatine concentration.

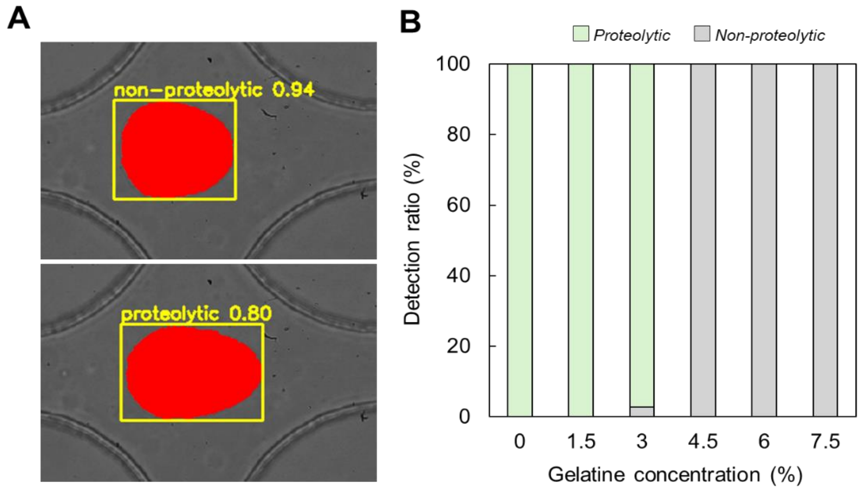


**Figure S3. Validation and confidence in gelatine droplet detection.** Automated detection of 7.5% and 0% gelatine droplets is shown within the ROI, identified via YOLO analysis (A) with tags indicating the detection confidence scores for proteolytic (liquid) and non-proteolytic (solid) droplet classes. The bar chart presents the results of experiments using six different gelatine concentrations, with over 4,500 droplets detected in total (B).

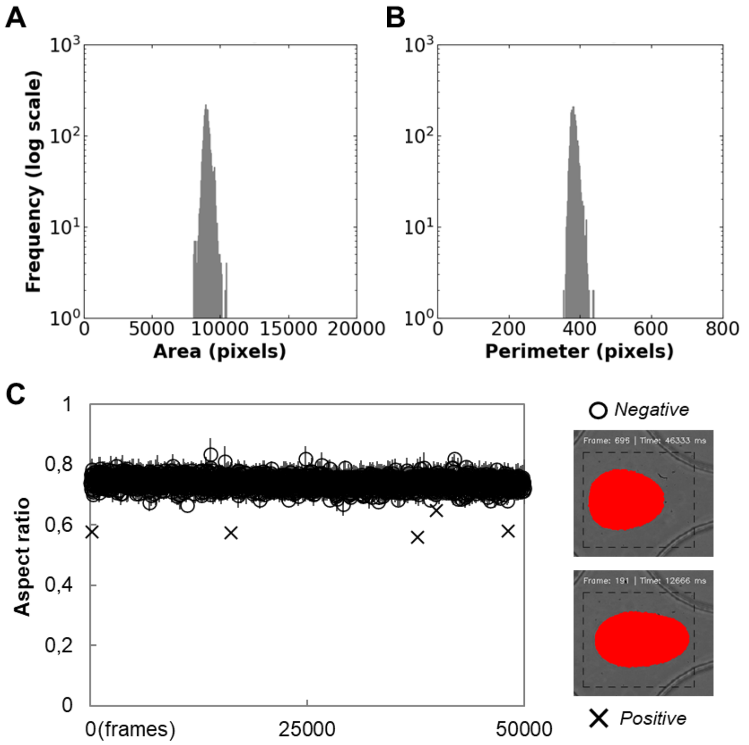


**Figure S4. IDO analysis of wastewater sludge.** The image-based method we propose represents a highly customisable system capable of adapting to different video conditions. Here, we report the Area (A) and Perimeter (B) thresholds set for droplet screening. These thresholds eliminate merged and split droplets from detection, enabling the exclusive characterisation of monodisperse droplets. The area threshold was set to include droplets ranging from 8000 to 10,500 pixels, accounting for the deformation observed in liquid droplets within the ROI during squeezing. Droplets obtained from an environmental screening predominantly exhibited an aspect ratio of approximately 0.75-0.8, whereas positive liquefied droplets were characterised by values <0.65 (C).

**Video S1. High viscosity gelatine droplets generation.** Emulsions were generated in a 50 x 50 µm flow-focusing device. The maximum throughput we achieved by using the final concentration of gelatine of 75 g/L to generate droplets (100 pL) was around 1.5-2 kHz.

**Video S2-S3. Screening of LB 0.5x and LB 0.5x + 7.5% gelatine droplets.** Droplets containing different concentrations of gelatine were analyzed within the ROI of the IDO device. This video illustrates the level of deformation detected for the minimum (LB0.5x) and maximum (LB0.5x + 7.5% gel) gelatine concentrations. The LB0.5x + 7.5% gelatine droplets were marked with 0.5% (v/v) polystyrene microbeads of approximately 1 µm in diameter.

**Video S4. Screening of mock consortia.** IDO analysis for the quantification of proteolytic strains was preliminarily assessed using a synthetic two-strain microbial community, composed of a highly proteolytic reference strain (*P. aeruginosa*) and a non-proteolytic strain (*E. coli*), clonally cultivated in droplets.

**Video S5-S6. Environmental screening of wastewater sludge.** The screening of microbial activity in a wastewater sludge sample was conducted after 24 hours of dynamic droplet incubation. Droplets containing empty or non-proteolytic strains maintained a spherical shape, whereas droplets with proteolytic activity became oval-shaped. The sorting frequency was approximately 0.25 kHz.

### References


1. Qin D, Xia Y, Whitesides GM. Soft lithography for micro- and nanoscale patterning. *Nature Protocols*. 2010;5(3):491-502. doi:10.1038/nprot.2009.234

2. Jenkins G. Rapid prototyping of PDMS devices using SU-8 lithography. *Microfluidic Diagnostics: Methods and Protocols*. Published online 2013:153-168.

3. Neun S, Kaminski TS, Hollfelder F. Chapter Five - Single-cell activity screening in microfluidic droplets. In: Allbritton NL, Kovarik ML, eds. *Methods in Enzymology*. Vol 628. Academic Press; 2019:95-112. doi:10.1016/bs.mie.2019.07.009

4. Mahler L, Tovar MA, Weber T, et al. Enhanced and homogeneous oxygen availability during incubation of microfluidic droplets. *RSC Adv*. 2015;5. doi:10.1039/C5RA20118G
